# Translation of cellular protein localization by generative adversarial network

**DOI:** 10.1101/2021.04.23.441224

**Authors:** Kei Shigene, Yuta Hiasa, Yoshito Otake, Mazen Soufi, Suphamon Janewanthanakul, Tamako Nishimura, Yoshinobu Sato, Shiro Suetsugu

## Abstract

The protein localization in cells had been analyzed by the fluorescent labeling by indirect immunofluorescence and fluorescent protein tagging. However, the relationships between the localizations between different proteins had not been analyzed by artificial intelligence. In this study, we applied the generative adversarial network (GAN) to generate the protein localizations each other, in which the generation was dependent on the types of cells and the relationships between the proteins. Lamellipodia are one of the actin-dependent subcellular structures involved in cell migration and are mainly generated by the Wiskott-Aldrich syndrome protein (WASP)-family verprolin homologous protein 2 (WAVE2) and the membrane remodeling I-BAR domain protein IRSp53. Focal adhesions are another actin-based structure that contains vinculin protein and are essential for cell migration. In contrast, microtubules are not thought to be directly related to actin filaments. The GAN was trained using images of actin filaments paired with WAVE2, vinculin, IRSp53, and microtubules. Then, the generated images of WAVE2, vinculin, and IRSp53 by the GAN showed high similarity to the real images of WAVE2, vinculin, and IRSp53, respectively. However, the microtubule images generated from actin filament images were inferior, corroborating that the microscopic images of actin filaments provide more information about actin-related protein localization. Collectively, this study suggests that the image translation by the GAN can predict the localization of functionally related proteins.

## Introduction

Machine learning has achieved significant success in various fields (Moen et al., 2019). Machine learning has been applied for classifying cellular images (Brent and Boucheron, 2018;Camacho et al., 2018;Moen et al., 2019). Using bright-field cell images, radiation-resistant cells were distinguished from the parental cells by machine learning (Toratani et al., 2018). Breast cancer cells treated with the anti-cancer agent paclitaxel were also distinguished from the non-treated ones by machine learning (Kobayashi et al., 2017). Furthermore, the direction of cell migration was predicted from sequences of cell images (Nishimoto et al., 2019). These results demonstrate that machine learning can extract information associated with cellular properties from images.

Machine learning has been applied not only in the classifications mentioned above, but also in protein localization. For example, a method known as *in silico* labeling reportedly generated a putative stained image of a specific marker protein from bright-field cell images to identify the nuclei, neural cells, and live cells (Christiansen et al., 2018). The automatic segmentation of intracellular organelles such as the Golgi apparatus and endoplasmic reticulum from bright-field cell images was also achieved (Pärnamaa and Parts, 2017). However, the translation of the localization of one molecule to another has not been reported.

The generative adversarial network (GAN) is one of the methods in image analysis by machine learning, where the probability distribution model obtained through training with a number of paired images generates hypothetical paired images (Goodfellow et al., 2014). Here, the GAN can generate a similar image of A from an image of B, after learning many paired images of A and B. For example, the GAN can reportedly generate an image of a “smiling” face from an image with a “non-smiling” face, by learning a lot of paired images of non-smiling and smiling faces (Sagawa and Hagiwara, 2018). The GAN comprises two components: a generator and a discriminator, and thus can generate high-quality images by the competition between the generator and the discriminator. Pix2pix is one of the major implementations of the GAN in image-to-image translation problems (Isola et al., 2017). Pix2pix successfully generated many kinds of paired images, including a map from an aerial image, a color image from a black-and-white image, a label to a street scene, a biomedical image like that of an MRI to the labels of the organs, and so on.

In cell biology, Pix2pix has been applied for labeling cellular membranes and nuclei, using images of these markers (Tsuda and Hotta, 2019). In this case, the label generation was performed by training with the image pairs of the labels in the regions of the membrane and nucleus (the label images) and the actual images. However, almost no report has demonstrated the application of pix2pix to the generation of an image showing the cellular molecule localization at subcellular resolutions, i.e., the generation of images showing the localization of a protein from those of other proteins. We hypothesized that pix2pix could be used to generate, i.e., to predict, the protein localization.

Cells change their shapes based on the mitotic cycle, surrounding environment, and various other situations by altering the cytoskeleton, including actin filaments (Pollard and Borisy, 2003;Gunning et al., 2015). In cells, actin filaments further assemble into higher-order configurations, which are primarily determined by Rho-family small GTPases, including Cdc42, RhoA, and Rac1 (Hall, 1998;Takai et al., 2001). Among them, Rac1 induces actin filament branching through WAVE2 (Bear et al., 1998;Machesky and Insall, 1998;Miki et al., 1998;Suetsugu et al., 1999b;Suetsugu et al., 2003). The activation of Rac1 induces conformational changes of WAVE2 in the regulatory complex, consisting of Sra1/PIR121, WAVE2, Nap1, Abi1/2, and HSPC300/BRICK, leading to the activation of the Arp2/3 complex within the branched actin filaments (Innocenti et al., 2004;Suetsugu et al., 2006;Ismail et al., 2009;Chen et al., 2010) (Figure 1A). IRSp53 is also involved in this lamellipodia formation, through WAVE2 (Miki et al., 2000;Suetsugu et al., 2006). Vinculin is a protein at focal adhesions, which are connected to actin filaments and are essential for lamellipodia formation (Ziegler et al., 2006). Lamellipodia are regarded as one of the essential structures for cell migration, including cancer cell invasion and metastasis (Takenawa and Suetsugu, 2007;Ridley, 2011). In this study, we successfully translated the images of actin filaments of cells to those of WAVE2, IRSp53, and vinculin, suggesting that the image translation by the GAN can generate the images of functionally related proteins.

## Results

### Prediction of WAVE2 localization from images of actin filaments by conditional GANs

We used Swiss 3T3 cells because they form lamellipodia upon the activation of Rac1 (Ridley et al., 1992). We introduced a constitutively active Rac1 mutant into Swiss 3T3 cells to induce lamellipodia. After chemical fixation, the cells were stained with phalloidin and an anti-WAVE2 antibody to visualize actin filaments and WAVE2, respectively. The fan-shaped actin filament substructures at the cell periphery, which were assumed to be lamellipodia, had WAVE2 (Figure 1B and C).

**Figure 1.**
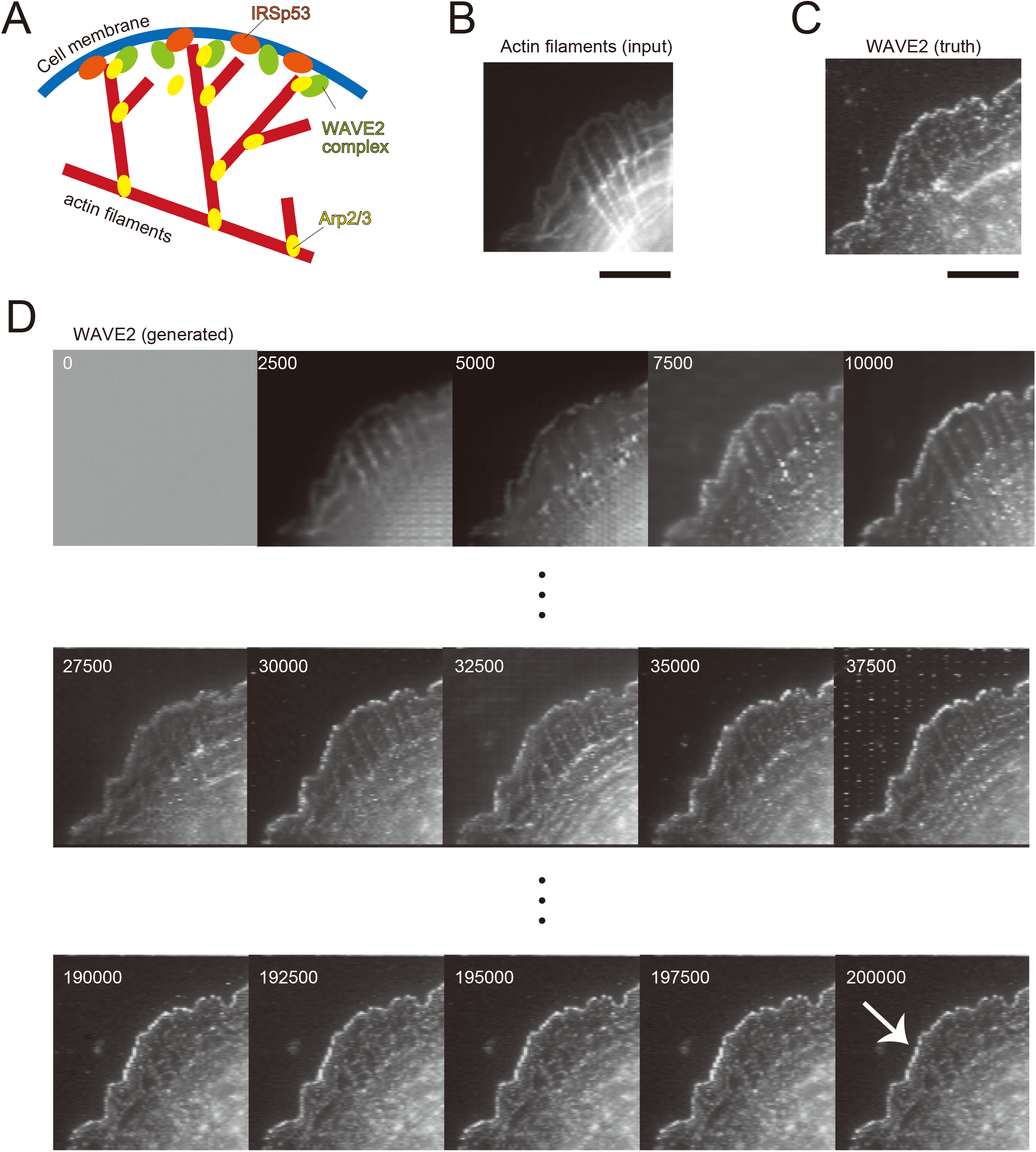
Lamellipodia and WAVE2 localization in Swiss 3T3 cells. (A) Schematic illustration of the configuration of actin filaments and WAVE2 localization at lamellipodia. Upon Rac1 activation, the WAVE2 in the protein complex is activated, leading to the activation of the Arp2/3 complex for branched actin filament formation. IRSp53 cooperates with WAVE2 for its activation by Rac1 at the plasma membrane. (B) Input image of actin filaments in Swiss 3T3 cells expressing the active form of Rac1. Actin filaments were stained by Rhodamine–phalloidin. Lamellipodia are fan-shaped structures formed at cell edges. (C) Actual WAVE2 image co-immunostained with (B), showing accumulation at the edges of lamellipodia. (D) Progress of the WAVE2 image generation. Images are shown at every 2,500 iterations (1 epoch). The iteration number is shown in the images. Image generation starts with a gray image without any features, as shown at iteration 0. The arrow indicates the lamellipodia. Scale bars, 10 µm.

Pairs of images of actin filaments and WAVE2 were taken and used for the training of the pix2pix conditional GAN model. The detailed methodology is described in the Conditional GAN subsection of the Material and Method section (Hiasa et al., 2019). The translation performance was estimated by four-fold cross-validation, with 772 paired images of actin filaments and WAVE2. In each subset, the training set comprised 579 images, of which 15% were used as the validation set. No augmentation was performed for the images of Swiss 3T3 cells. The remaining 193 images were used for the testing set. The number of iterations, which corresponds to the epoch number for the training, was 200,000. This process was repeated four times. As the number of iterations grew, the similarity between the generated and actual WAVE2 images increased (Figure 1D). The generated WAVE2 final images were similar to those obtained by antibody staining.

Examples of the generated WAVE2 images are presented in Figure 2. WAVE2 showed the prominent localization to lamellipodia. WAVE2 localization was generated clearly at the edge of the cells (Figure 2A-C). The pix2pix model predicted the localization of WAVE2 regardless of the size of the lamellipodia (Figure 2A-C). WAVE2 is not only localized at lamellipodia, but also at other subcellular structures of actin filaments. The tips of microspikes or filopodia in the lamellipodia are known to have a WAVE2 distribution (Figure 2A) (Nakagawa et al., 2003;Nozumi et al., 2003). The pix2pix model could predict the WAVE2 localization at the microspike structures in lamellipodia, as indicated by the dashed square in Figure 2A. Interestingly, the microspikes outside of the lamellipodia were also predicted to have WAVE2, as indicated by the square in Figure 2A, and indeed had real WAVE2. WAVE2 also reportedly functions at the cell-cell junctions (Yamazaki et al., 2007;Nishimura et al., 2016). The two cells were in contact with each other, with the WAVE2 localization at the contact sites (Figure 2D). WAVE2 localization was generated clearly between the cell-cell contacts (Figure 2D). The overall predicted WAVE2 localization by pix2pix appeared to be quite similar to the real WAVE2 localization detected by antibody staining. Together, these facts suggest that pix2pix can predict the localization of WAVE2 not only in the lamellipodia but also in other cellular structures.

**Figure 2.**
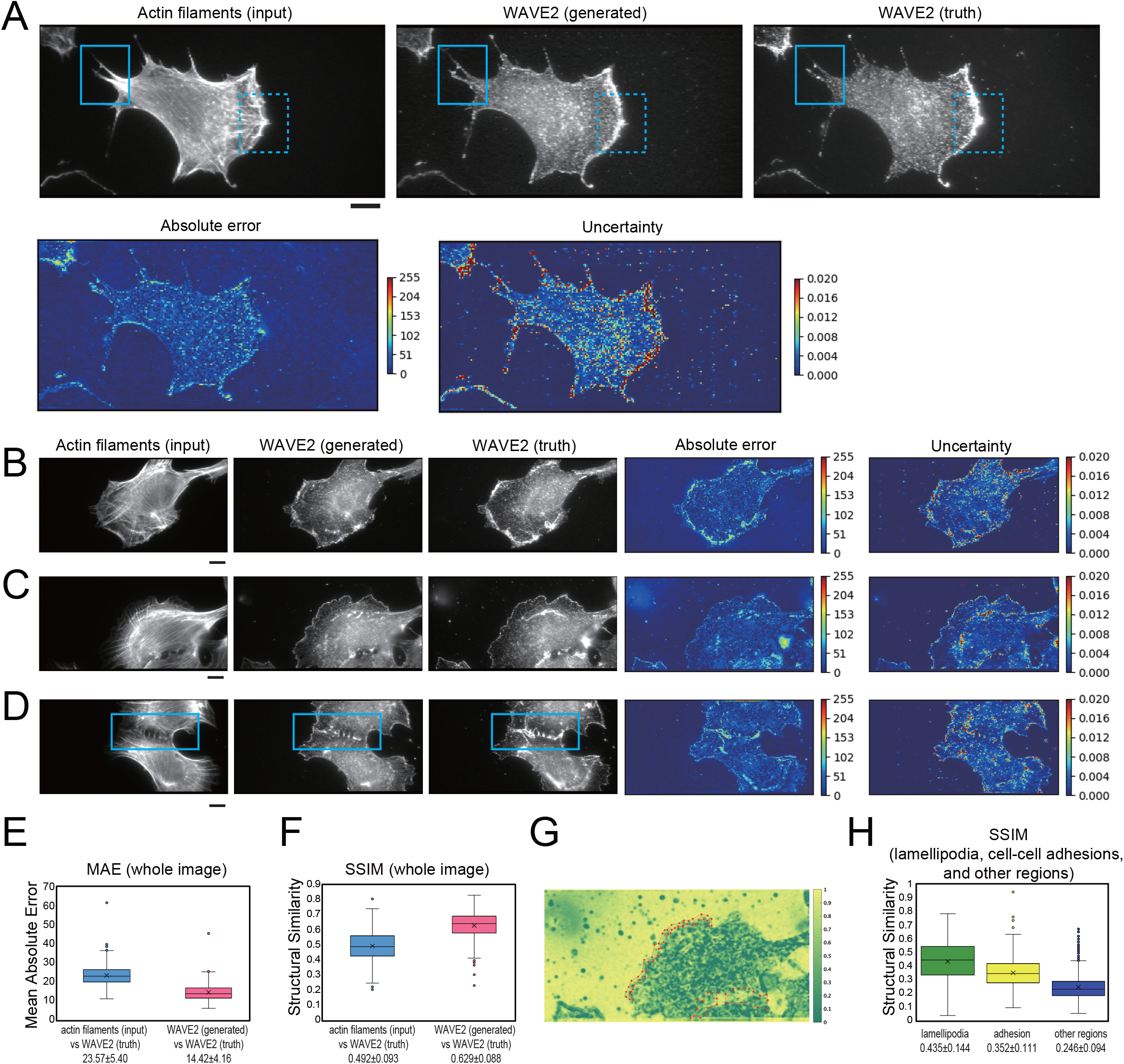
Generation of WAVE2 images from actin filaments in Swiss 3T3 cells. (A) Generation of a WAVE2 image by pix2pix from an actin-filament image. The cells were stained with phalloidin for actin filaments and with an anti-WAVE2 antibody after fixation and permeabilization. An input image (actin filament image), an output image (generated WAVE2 image), a ground truth image (WAVE2 immuno-stained image), an absolute error image, and an uncertainty image are shown. The microspikes in the lamellipodia are marked with dashed squares, and those outside of the lamellipodia are marked with solid squares. Absolute error represents the difference in WAVE2 values in each pixel. Uncertainty in image generation represents fluctuation of WAVE2 values based on various “dropouts” in convolutional neural networks, i.e., the robustness of the generation in each pixel. With higher values of absolute errors and uncertainty, the color of the heat map becomes closer to red. Scale bars, 10 µm. (B, C) Image generation of WAVE2 images of cells with various sizes of lamellipodia, as shown in (A). Scale bars, 10 µm. (D) Generation of a WAVE2 image of cells that formed a cell-cell adhesion marked by a rectangle. Scale bars, 10 µm. (E) Box plot of the mean absolute error (MAE) between the generated and actual WAVE2 images, as well as between images of actin filaments (input) and actual WAVE2 as a reference. Quantification was performed for the entire images with four-fold cross-validation (n=772). The mean values are shown at the bottom. (F) Box plot of the structural similarity index measure (SSIM) value between the generated and actual WAVE2 images, as well as between images of actin filaments and actual WAVE2 as a reference. Quantification was performed for all images with four-fold cross-validation (n=772). The mean values are shown at the bottom. (G) Representative SSIM map corresponding to the image in Figure 2C, showing the structural similarity at each 11 × 11 pixel window. Regions of lamellipodia are marked with polygons. (H) Box plot of the SSIM values from 1,926 pairs of lamellipodia, cell-cell adhesions, and other cellular regions.

### Performance of the prediction of WAVE2 localization

We evaluated the prediction accuracy in each pixel. The absolute error in Figure 2A-D is the difference between the generated and true WAVE2 in each pixel, which at lamellipodia was higher than that at the background (Figure 2A-D).

Another estimation of the accuracy is based on the robustness of the prediction. The uncertainty in this translation had been difficult to describe. However, the uncertainty of such labeling was successfully demonstrated using Bayesian convolutional neural networks (Bayesian CNNs), based on the U-net architecture combined with the Monte-Carlo dropout of the network layers (Hiasa et al., 2019). The dropout (removal) of the network layers resulted in different outputs; however, the high probability output contains less dependency on the alteration of the network layers, resulting in less uncertainty in the output. The uncertainty in predictions of WAVE2 localization was also high at the lamellipodia (Figure 2A-D).

The absolute error and the uncertainty of the prediction were typically higher at the pixels of WAVE2 localization, indicating that the intensity of WAVE2 localization is not predicted in the absolute values at a pixel resolution; instead, the prediction is more qualitative, reflecting the context of actin filaments for WAVE2 localization. Therefore, the absolute error would be caused by the aleatoric uncertainty from the randomness of the measurements, rather than the epistemic uncertainty of the prediction.

The overall image prediction was then estimated by the mean absolute error (MAE) and the structural similarity index measurement (SSIM) between the generated and true WAVE2 images. MAE is the mean absolute difference in the pixel values, which is related to the absolute errors in each image. Therefore, a smaller MAE indicates a higher similarity between the two images. SSIM is based on the variance in the pixel values. Therefore, higher SSIM indicates more similarity in the perceived quality. The MAE between the generated and true images was lower, and the SSIM between the generated and true images was higher, than those between the true actin filament (input) and true WAVE2 images, suggesting that the generator successfully produced WAVE2 images that were better representations than the input actin filament images (Figure 2E and F).

Subsequently, we analyzed the performance of WAVE2 localization prediction at the subcellular level. The SSIM was calculated for each 11 × 11 pixel window to generate the SSIM map, and the representative analysis corresponding to Figure 2C is shown in Figure 2G. Then, the images of the ground truth were manually annotated using Labelme (Russell et al., 2008) (Figure 2G), which enclosed parts of the image with polygons and saved the region information in the JSON file format. The SSIM of the manually annotated lamellipodia and cell-cell adhesions, where WAVE2 was localized, were compared with the SSIM in the other cellular regions. The average SSIMs of lamellipodia and cell–cell adhesion sites were higher than the average SSIMs in the non-lamellipodia regions (Figure 2H). This shows the possibility of using GAN in quantifying the lamellipodia formation.

### Application to the lifeact staining of actin filaments and IRSp53

To examine the generalization of this method, we trained the pix2pix model using glioma U251 cells and another regulator of WAVE2, IRSp53 (Miki et al., 2000;Suetsugu et al., 2006). The IRSp53-knockout U251 cells were prepared, and the IRSp53 expression was restored by the stable expression of GFP-IRSp53. The cells were then further stably labeled by lifeact tagged with mCherry (lifeact-mCherry) to visualize actin filaments (Riedl et al., 2008). These U251 cells were cultured in serum and fixed for observations of the lamellipodia without the forced activation of Rac1. The WAVE2 localization was identified by the antibody staining. The images of the actin filaments by lifeact, IRSp53 by GFP, and WAVE2 by the antibody staining were subjected to the machine learning for image translation using pix2pix. We attempted to translate the images of actin filaments (lifeact) to WAVE2, actin filaments to IRSp53, and IRSp53 to WAVE2. The translation performance was estimated by four-fold cross-validation. In total, 100 images were obtained, and 75 were subjected to the training. The 75 images were augmented 7-fold by rotations at 90-degree steps and vertical and horizontal flipping, resulting in the training data set composed of 525 images, of which 15% were used as the validation set. The remaining 25 images were used for the testing set. The results showed that a lifeact image could produce the images of WAVE2 and IRSp53 (Figure 3A, B). Furthermore, pix2pix translated the IRSp53 images to the WAVE2 images (Figure 3C). The prediction at lamellipodia was regarded as having good quality, because the MAE values between the truth and generated images were lower than the MAE values between the input and truth images (Figure 3D). However, some features were not predicted, such as the staining around the nucleus (Figure 3A-B). The failure in the prediction of nuclear localization appears to reduce the quality in perception, because the SSIM values between the input and truth images were similar to those between the truth and generated images (Figure 3E).

**Figure 3.**
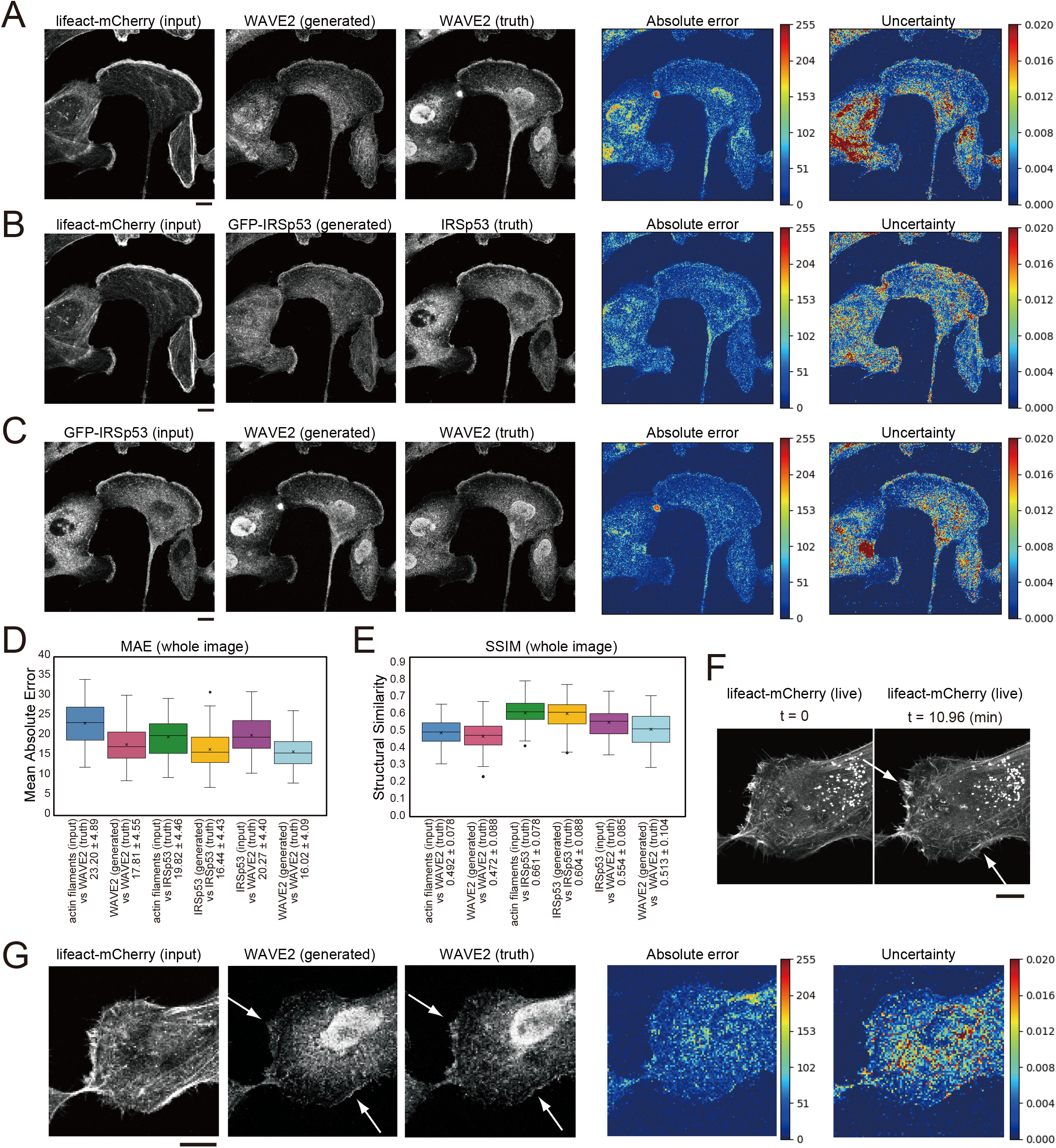
Generation of WAVE2 and IRSp53 images from actin filaments and WAVE2 images in IRSp53-expressing U251 cells. (A) Generation of a WAVE2 image from an actin filament image, by lifeact. The IRSp53-knocked-out U251 cells re-expressing GFP-tagged IRSp53 and lifeact-mCherry were stained with an anti-WAVE2 antibody after fixation and permeabilization. The results are depicted as in Figure 2A. Scale bar, 10 µm. (B) Generation of an IRSp53 image from an actin filament-stained image. Scale bar, 10 µm. (C) Image generation into a WAVE2 image from an IRSp53-stained image. Scale bar, 10 µm. (D) Box plot of the MAE of the entire images in four-fold cross-validation (n=100). (E) Box plot of the SSIM of the entire images in four-fold cross-validation (n=100). (F) Lamellipodia structures (arrows) observed in live imaging of lifeact-mCherry. Scale bar, 10 µm. (G) Generation of the WAVE2 image from the lifeact-mCherry image. The cells observed in (F) were fixed, permeabilized, and immunostained for WAVE2 (truth). The arrows indicate lamellipodia. Scale bars, 10 µm.

We observed the lifeact-mCherry in live cells to identify the lamellipodia at the leading edge (Figure 3F). The cells were then fixed, permeabilized, and stained for the WAVE2 localization. The permeabilization slightly altered the lifeact images, because the free lifeact in the cytosol was probably removed by permeabilization. The active lamellipodia region was stained with WAVE2, and the actin filament images for these lamellipodia were able to generate the WAVE2 image by using the trained model as above (Figure 3G). From these results, we conclude that pix2pix can specifically predict WAVE2 localization at the leading edge of lamellipodia under different conditions.

### Application to vinculin and tubulin localization

To examine whether GAN can be applicable to other molecules that are related to lamellipodia, we trained the model between actin filaments and vinculin staining. The translation performance of the model, trained by 100-paired actin filaments and vinculin images of U251 cells, was estimated by four-fold cross-validation. The 75 images were augmented 7-fold, resulting in the training data set composed of 525 images, of which 15% were used as the validation set. The remaining 25 images were used for the testing set. The pix2pix succeeded in generating vinculin images from actin-filament images (Figure 4A). The prediction of vinculin was regarded as having good quality, as judged by the MAE and the SSIM values (Figure 4B, C), as well as by the recognition by human eyes.

**Figure 4.**
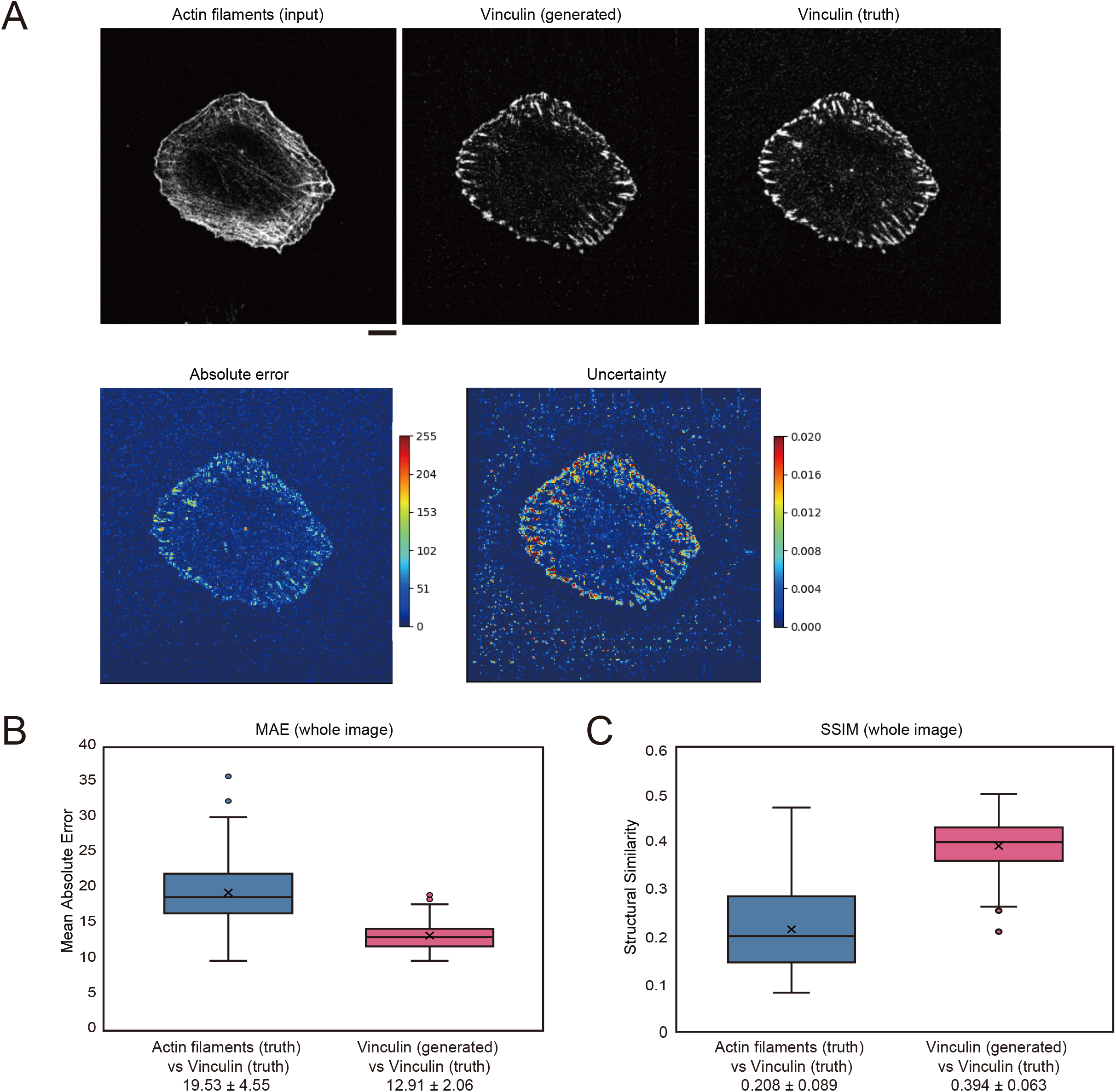
Generation of vinculin and tubulin images in U251 cells. (A) Generation of a vinculin image from an actin filament-stained image. U251 cells were stained with phalloidin for actin filaments and with an anti-vinculin antibody after fixation and permeabilization. The results are depicted as in Figure 2A. Scale bar, 10 µm. (B) Box plot of the MAE of entire images in four-fold cross-validation for (A) (n=100). (C) Box plot of the SSIM of entire images in four-fold cross-validation for (A) (n=100).

### Application to tubulin localization

To examine whether GAN can be applicable to other molecules that are not strongly related to actin filaments, we trained the model between actin filaments and tubulin staining of U251 cells as for the vinculin analysis. The pix2pix model generated a tubulin-like image from actin-filament images (Figure 5). However, the generated image did not reflect the feature of radial-like tubulin distribution (Figure 5). From these results, we conclude that the accuracy of pix2pix is decreased if the two molecules lack a strong correlation.

**Figure 5.**
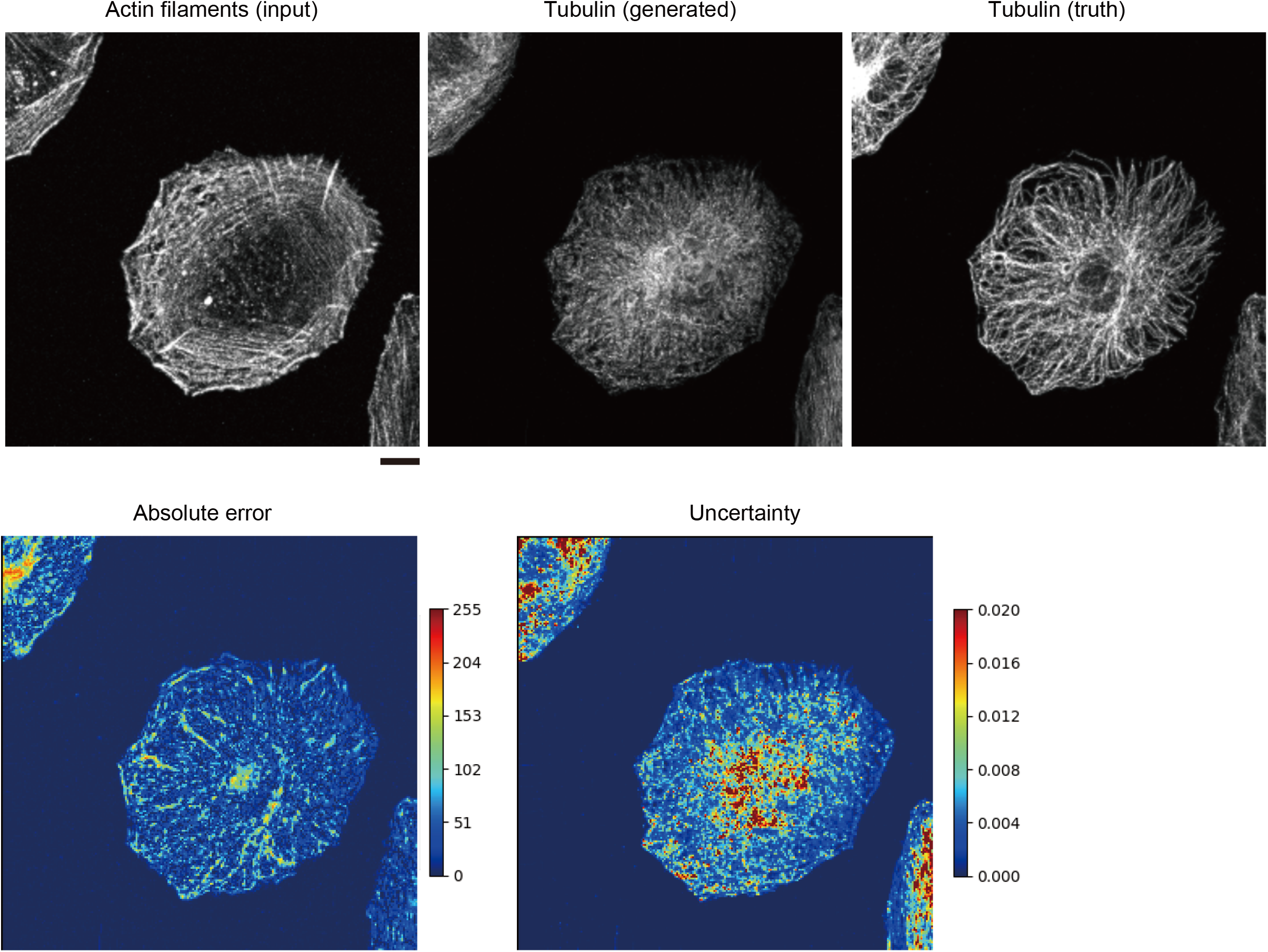
Generation of tubulin images in U251 cells. Generation of a tubulin image from an actin filament-stained image. U251 cells were stained with phalloidin for actin filaments and with an anti-*α*-tubulin antibody after fixation and permeabilization. The results are depicted as in Figure 2A. Scale bar, 10 µm.

## Discussion

In this study, we identified subcellular structures using a machine learning method, pix2pix-mediated image translation. Pix2pix is a GAN implementation, and WAVE2, IRSp53, and vinculin are established regulators of lamellipodia. Therefore, the prediction of these localizations by image translation is suggested to annotate lamellipodia among actin cytoskeletal structures. The experts in the field will easily distinguish lamellipodial actin filaments from non-lamellipodial ones, but sometimes lamellipodia are not obvious to the untrained eye. The judgment criteria for microscopic images will differ between researchers in different labs. At the very least, the GAN method will reproduce comparable results if the image qualities are similar. Furthermore, this method can be used to label lamellipodia in live cells to quantify the degree of lamellipodia formation, if computation speed increases in the future.

The prediction of WAVE2 localization was independent of the size of lamellipodia (Figure 2). The kernel size of the algorithm, which uses the 4-pixel window, is equivalent to ∼1.9 μm^2^. Therefore, the continuous features above this size are expected to be difficult to be predicted. However, the features of actin filaments at lamellipodia are thought to be within this window, and thus various sizes of lamellipodia could be annotated by the predicted WAVE2 localization. The vinculin and IRSp53 localizations are also thought to be predictable with such features of actin filaments in 4-pixel window.

We tried to predict the microtubule localization from actin filament localization; however, this predicted localization was not filamentous. These non-continuous filaments might arise from the kernel size. Alternatively, microtubules are not directly linked to actin filaments, in contrast to the regulators of actin filaments like WAVE2, IRSp53, and vinculin, resulting in the inaccurate prediction. In this respect, the GAN might be able to determine/predict the hierarchy of various proteins for specific cellular structures.

If the difference between the experimental and predicted images resulted from the mutually independent localization and function, then the GAN might be used as a tool to discover a localization and a function that can be independent of each other. The failure of the prediction of the nuclear staining of WAVE2 in U251 cells might indeed suggest the actin filament-independent function of WAVE2 (Figure 3). The nuclear staining of WAVE2 might represent either non-specific staining or the nuclear localization of WAVE2, as the nuclear localization of WAVE1 has been reported (Miyamoto and Gurdon, 2013). Interestingly, such nuclear staining of WAVE2 was not observed in Swiss 3T3 cells, suggesting that nuclear WAVE2 in U251 cells could be a result of non-specific staining (Figure 2). Therefore, the prediction is thought to require training dependent on the cell types, and might reflect the specific features of the cells. These inconsistencies in WAVE2 localizations between the generated and true images therefore might be due to insufficient learning, the randomness in experimental errors, or the true difference in the localization and functions that resulted from the independence of the observed pair of molecules. Further investigations will be required to prove these ideas; however, image correlation using the GAN might be performed to investigate the known and unknown spatial relations of paired molecules using localization images.

The prediction of protein localization could have great potential. The labeling by antibodies is normally limited to several proteins. In contrast, artificial staining can predict unlimited number of protein localizations from a single staining, which would be useful to detect the relationships between many molecules. The prediction should be complemented with the actual experiments. However, the prediction of molecule localizations in cells would at least be useful to search for the direction of study when dissecting the localization-related hierarchy of the proteins and their functions.

## Material and Method

### Cell culture

Plat-E, Swiss 3T3, and U251 cells were cultured in Dulbecco’s Modified Eagle Medium (DMEM) (Nacalai Tesque, 08459-64), supplemented with 10% fetal calf serum (FCS) and penicillin–streptomycin (PS) (DMEM-10% FCS/PS), at 37 °C in a 5% CO_2_ incubator. The Plat-E, Swiss 3T3, and A549 cells were passaged every 4, 3, and 2 d, respectively.

### Retrovirus-mediated gene transfer

Swiss 3T3 cells were transfected using the pMX-Myc-Rac1-CA vector (Suetsugu et al., 1999a), as follows. First, Plat-E cells were cultured overnight in a 12-well plate in DMEM-10% FCS/PS. For gene transfer, 100 µL of Opti-MEM with 1.6 µg of vector and 100 µL of Opti-MEM with 1 µL of 293 fectin transfection reagents (Thermo Fisher) were mixed, allowed to form a complex at room temperature for 20 min, and then added to the Plat-E cells in 0.8 ml medium (Kitamura et al., 2003). After 48 h, the culture supernatant was filtered using a 0.22 µm filter, and added to the cells in 1.2 ml medium with polybrene, at a concentration of 8 µg/mL. After 24 h, the medium was replaced with fresh DMEM-10% FCS/PS. After an additional 24 h, the cells were replated on a 24-well plate containing a coverslip (Matsunami) and cultured for another 48 h.

The IRSp53-knockout U251 cell line expressing GFP-IRSp53 was established using CRISPR/Cas9-mediated genome editing. The guide RNA targeting the second exon (29^th^ amino-acid residue) of IRSp53 (CCATGGCGATGAAGTTCCGG) was designed using the server http://crispr.mit.edu (Hsu et al., 2013) and inserted into the pX330 vector, which was transfected into the cells and then cloned (Mashiko et al., 2013). The expression of GFP-IRSp53 and lifeact-mCherry was performed by using the retrovirus as above, and then clones were isolated by using a fluorescence-activated cell sorter.

### Immunofluorescent staining of Swiss 3T3 and U251 cells

The cells were fixed with 4% paraformaldehyde in PBS for 20 min at room temperature. Subsequently, they were permeabilized with 0.5% Triton X-100 in PBS for 20 min at room temperature with gentle shaking. The cells were then washed with 0.1% Triton X-100 in PBS (PBS-T). Next, PBS containing 3% bovine serum albumin and 10% goat serum was added to block the cells for 1 h with gentle shaking. The cells were then washed with PBS-T. The primary antibody, either a rabbit anti-WAVE2 antibody (Cell Signaling, # 3659S), a mouse anti-vinculin (SIGMA, V 9131), a mouse anti-alpha-tubulin (SIGMA, clone DM1A) was diluted 100-fold, 200-fold, and 500-fold respectively, in the blocking solution, incubated for 1 h with gentle shaking, and then washed three times with PBS-T. The secondary antibody, an Alexa Fluor 488-goat anti-rabbit or mouse IgG antibody (highly cross-absorbed, Thermo-Fisher) diluted 400-fold, and rhodamine-phalloidin (Thermo-Fisher) for actin filament detection, diluted 1,000-fold in the blocking solution, were added and then incubated for 1 h with gentle shaking in the dark. The cells were then washed with PBS-T and mounted on a glass slide, using Prolong Diamond Antifade Mountant with DAPI (Thermo Fisher), allowed to solidify at room temperature overnight, and then stored at 4 °C. Swiss 3T3 cells were observed using an IX81 fluorescence microscope (OLYMPUS) with W-View Gemini (Hamamatsu Photonics). U251 cells and A549 cells were observed using an FV1000 confocal microscope (Olympus).

### Conditional GANs

The purpose of this study was to determine the conditional distribution of WAVE2 based on actin filaments. The pix2pix conditional GAN (Isola et al., 2017), allows the patch discriminator to capture the Markov property of the image as an adversarial loss, allowing the transformed image to maintain high spatial frequencies. The formula for this adversarial learning is as follows: 

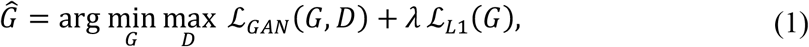

where a generator *G* translates images of actin filaments *x* to WAVE2 images *y*, which are trained to translate images of actin filaments that a discriminator *D* cannot distinguish from the “real” WAVE2 images by antibody staining, as follows: 

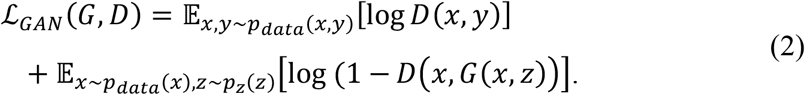

In addition to the adversarial loss, the conditional loss, which is the similarity between the “fake” and “real” WAVE2 images, is introduced as follows: 

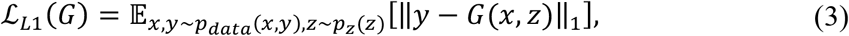

where *z* denotes the random noise.

We primarily followed this framework and extended the generator and discriminator networks. In this study, the generator was replaced with a Bayesian U-Net (Hiasa et al., 2019) for the uncertainty estimation. Spectral normalization (Miyato et al., 2018) was applied to the patch discriminator for stabilizing the optimization. In the inference phase, the predictive distribution is expressed as 

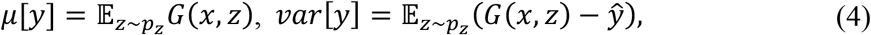

where *<u>μ</u>* and *var* denote the mean and variance, respectively.

WAVE2 and actin filament images were downscaled to 256 × 256 pixels and normalized such that the intensity of the [1st, 99th] percentile was mapped to [-1, 1]. Data augmentation was applied based on spatial transforms, including the translation of [−10, +10]% of the image size, rotation of [−10, +10]°, scale of [−10, +10]%, shear transformation with the shear angle of [−π/16, +π/16] rad, and flipping in the horizontal and vertical directions. The kernel size was 4 pixels, i.e., ∼1.9 μm^2^. The codes that were used, including the details of each network and training manner, are available at https://github.com/yuta-hi/bayesian_unet.

### Estimation of errors

The results were evaluated based on the MAE and SSIM. The MAE shows the absolute error in the brightness value in each pixel and is expressed as 

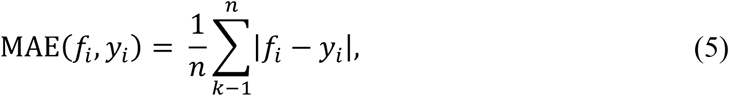

where *f*_*i*_ and *y*_*i*_ denote the true and predicted values, respectively.

The SSIM indicates the similarity of the average, variance, and covariance of surrounding pixels in terms of the brightness, contrast, and structure. Thus, it is an index that incorporates the correlation not only with individual pixels but also with the surrounding pixels. The SSIM is expressed as 

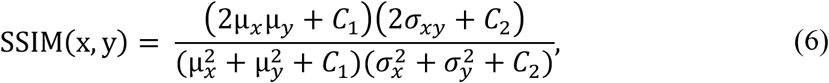

where x and y are the ground-truth (WAVE2) and predicted images, respectively, µ is the average pixel value, σ is the standard deviation of the pixel value, σ_xy_ is the covariance between x and y, C1 = (0.01 × L2), C2 = (0.03 × L2), and L is the dynamic range of the images (Wang et al., 2004). In this study, we used 8-bit images; hence, L = 255.

The lamellipodia regions were manually annotated for SSIM calculations using Labelme (Russell et al., 2008) (https://github.com/wkentaro/labelme) to extract the SSIM values at the lamellipodia.

## Data Availability Statement

The codes and datasets generated for this study are available upon request to the corresponding authors.

## Author contributions

KS, SJ, and TN performed the experiments. KS, YH, YO, SJ, and MS performed the computations. KS, YH, YO, MS, SJ, TN, YS, and SS analyzed the results, created the figures, and wrote the manuscript. YS and SS supervised the project.

## Acknowledgment

We thank all the laboratory members for their helpful discussions and continuous support for this study. We thank Prof. Naoki Watanabe at Kyoto University for lifeact.

